# Degradation Bottlenecks and Resource Competition in Transiently and Stably Engineered Mammalian Cells

**DOI:** 10.1101/2024.06.03.597247

**Authors:** Jacopo Gabrielli, Roberto Di Blasi, Cleo Kontoravdi, Francesca Ceroni

## Abstract

Degradation tags, otherwise known as degrons, are portable sequences that can be used to alter protein stability. Here, we report that degron-tagged proteins compete for cellular degradation resources in engineered mammalian cells leading to coupling of the degradation rates of otherwise independently expressed proteins when constitutively targeted human degrons are adopted. By adopting inducible bacterial and plant degrons we also highlight how orthogonality and uncoupling of synthetic construct degradation from the native machinery can be achieved. We show the effect of this competition to be dependent on the context of the degrons where C-terminal degradation appears to impact competition the most across our tested settings. We then build a genomically integrated capacity monitor tagged with different degrons and confirm resource competition between genomic and transiently expressed DNA constructs. This work expands the characterisation of resource competition in engineered mammalian cells to degradation also including integrated systems, providing a framework for the optimisation of heterologous expression systems to advance applications in fundamental and applied biological research.

## Introduction

Protein degradation is a physiological process conserved across organisms that regulates cellular protein homeostasis^1–4^. In mammalian cells, degradation occurs primarily through the ubiquitin-proteasome system (UPS). This involves three subsequent enzymatic steps mediated by the ubiquitin-activating enzyme E1, the ubiquitin-conjugating enzyme E2, and ubiquitin-protein ligase E3, respectively, which facilitate protein poly-ubiquitination and subsequent proteasomal degradation. E1 mediates the binding of ubiquitin through an ATP-dependent reaction, subsequently transferring it to E2. E3 then catalyses the transfer of ubiquitin onto the target protein by formation of an isopeptide bond. By repeated rounds of ubiquitination (i.e. poly-ubiquitination) the target proteins are directed to degradation through the proteasomal pathway^1,5–7^.

The likelihood for a protein to be degraded by the UPS can be enhanced by the addition of a sequence of amino acids (i.e. degron). Degrons can be present within the protein sequence, at its C-terminus or at the N-terminus. C-terminal and N-terminal degrons have the advantage to be portable from one sequence to another, thus enabling controlled degradation of any protein of interest^8–11^.

In synthetic biology, the number of proteins produced by engineered synthetic constructs and their degradation rate can drastically affect the functioning of gene and protein circuits. For this reason, degrons have gained popularity, with a variety of sequences developed as tools to achieve post-translational regulation of protein expression^12–15^. Among the applications, degrons have been used to accelerate transitions between gene circuit steady states^16–19^ and to achieve desired dynamic expression patterns^20^. While some degron sequences mediate constitutive protein targeting by the degradation machinery (constitutive degrons), inducible degrons that enable control of protein degradation upon addition of a specific inducer^15,21–26^, have also been adopted, with applications in the regulation of chimeric antigen receptor activity and cell therapies^27–29^.

One aspect that is usually overlooked in studies aimed at the design and characterisation of synthetic degradation systems, is their impact on the cellular pool of degradation resources. In bacteria, Hasty *et al*. showed how using multiple degrons that share part of the degradation pathway, in their case the ClpXP protease, led to coupling of degradation rates and enzymatic queueing^30,31^. Mather *et al*. expanded the concept of competition for degradation resources in bacteria by characterising crosstalk involving other bacterial proteases such as ClpAP and Lon^32,33^. Previous work on resource competition in mammalian cells characterised bottlenecks in the transcriptional, translational and secretory machinery during transient gene expression^34–38^. However, considerations of how limitations in protein degradation resources impact heterologous gene expression have not been reported for these systems, yet^39^.

Here we developed a framework for the characterisation of resource competition at the degradation level in mammalian cells, featuring both transient and stably integrated systems.

We started by selecting a small library of degrons from a recent study by Chassin *et al*. that reported the comprehensive characterisation of a library of degrons in mammalian cells^17^. Three classes of degrons were considered. First, two classes featuring an N-terminal ubiquitin protein, whose N-terminal fusion is known to mediate proteasomal degradation, bearing either an intact or a mutated isopeptidase site. An intact isopeptidase site leads to recognition by deubiquitinating enzymes, which cleave ubiquitin, revealing either a stabilizing or destabilizing amino acid at the N-terminus of the target protein to which the N-end rule applies^10^. Conversely, degradation domains with a mutated isopeptidase site evade recognition by isopeptidase enzymes targeting the protein to the UFD (Ub-Fusion Degradation)^40^. Both categories of degradation domains lead to degradation via the proteasome-mediated pathway^17^.The third class included three PEST (proline, glutamate, serine, threonine) degrons, posited to be independent of the ubiquitin proteasome system^41–44^.

Our selected library included i) two N-terminal ubiquitin tags featuring mutated isopeptidase sites, UbVR with a higher degradation rate, and 2xUbAV with two mutated ubiquitin domains and a lower degradation rate, ii) two N-terminal ubiquitin tags with intact isopeptidase sites, one with an N-terminal arginine and higher degradation rate, UbR, and one with an N-terminal methionine and a lower degradation rate, UbM, iii) two C-terminal PEST-based degrons, MODCPEST with a higher degradation rate, and PEST with a lower degradation rate.

We used these degrons to generate a library of EGFP expressing test constructs encompassing C-terminal, N-terminal degradation aiming at characterising differences in resource availability between the C-terminal and N-terminal degradation pathways. We also tested two widely adopted bacterial and plant degrons which have displayed high orthogonality in resource usage, making them amenable to synthetic systems with multiple degron-tagged proteins. Subsequently, we constructed and characterised a genomically integrated, inducible, mammalian capacity monitor and employed this to expand considerations of resource competition to genomically expressed constructs. Our study provides characterisation of competition for degradation resources in mammalian cells and suggests routes to achieve uncoupling between transiently expressed systems and the cellular degradation machinery to improve the engineering of heterologous genetic systems with desired behaviour. The capacity monitor cell line developed here can find applicability as a proxy measure for impact of synthetic gene circuits on native gene expression and protein degradation.

## Results

### Competition for degradation resources impacts mammalian expression systems

Capacity monitors have been previously adopted to quantify and visualise gene expression resource competition in bacteria and mammalian cells^34,35,45–48^. Monitors are fluorescent expression cassettes that function as proxy for the available gene expression capacity within a cellular host. When adopted in combination with co-expressed test constructs, a change in the monitor`s expression level can be considered a semi-quantitative proxy measure of the resource footprint of that given construct.

To investigate whether competition for degradation resources also impacts mammalian heterologous expression, we devised an experiment in which a PEST-tagged capacity monitor, designed by incorporating the PEST degradation domains into the fluorescent readout of transiently expressed monitors (mKATE), was co-transfected either with a MODCPEST-tagged or an untagged EGFP-expressing test construct (Figure 1A and B). Both constructs shared identical plasmid backbones, promoters, Kozak sequences, and polyA signals. The sole distinction between them lay in the presence or absence of the MODCPEST domain.

**Figure 1.**
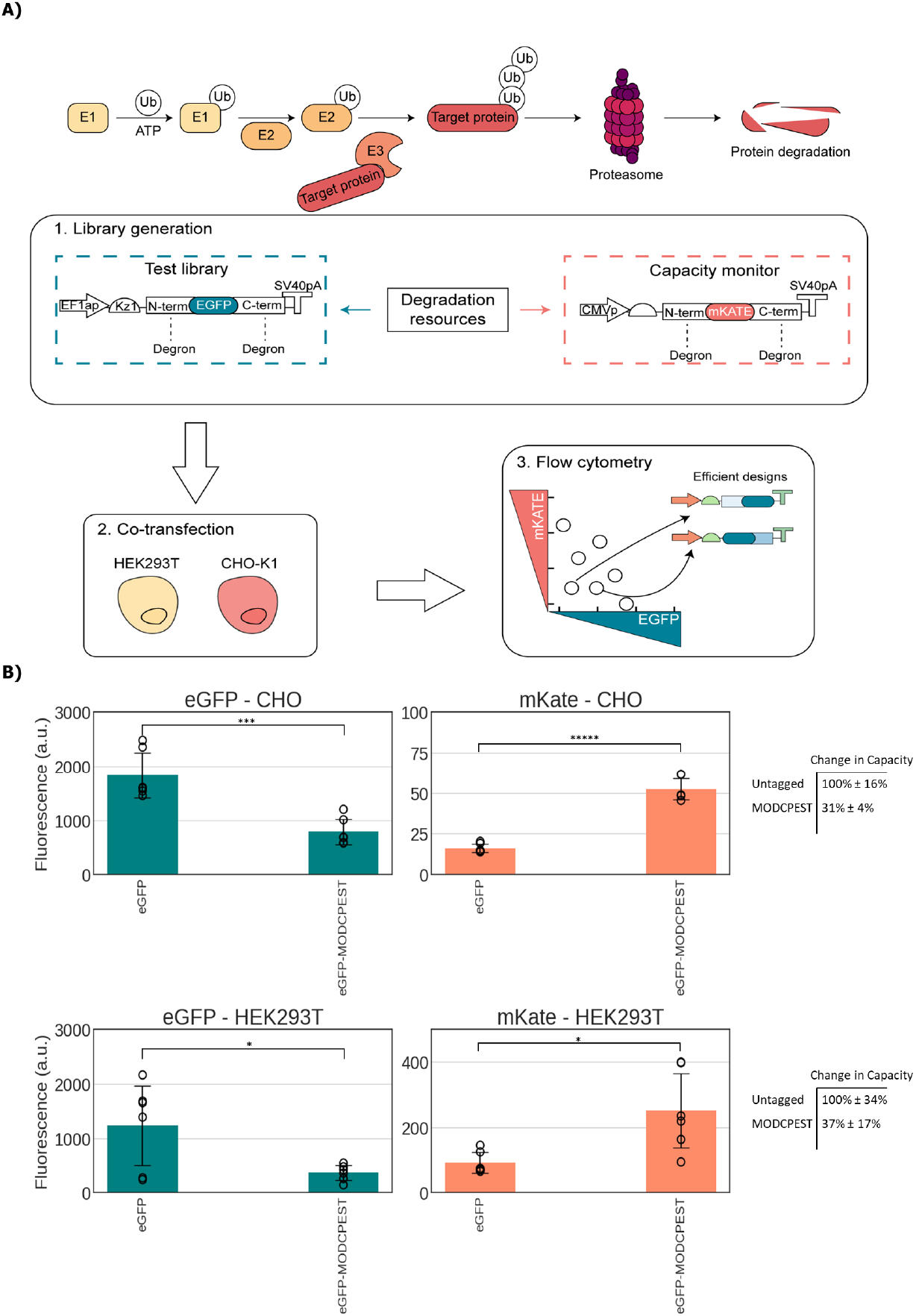
Probing degradation-induced coupling via capacity monitors. **(A)** Simplified diagram of the UPS degradation pathway followed by experimental workflow schematics. A library of test constructs, tagged with degrons, is co-transfected into HEK293T and CHO-K1 cell lines along with a destabilized capacity monitor. Intracellular protein levels are quantified using flow cytometry. The competition for degradation resources can be visualized by plotting the levels of the capacity monitor protein against the levels of the test constructs’ protein. In this plot, the constructs located at the bottom left corner represent the most efficient candidates. **(B)** Proof-of-concept experiment where a destabilized monitor, with a PEST degron, is co-transfected with a test construct tagged with the MODCPEST degron in HEK293T and CHO-K1 cells. Test plasmid and capacity monitor intracellular protein levels are reported as mean fluorescence (arbitrary units) ± std. The table reports 1/(mKATE of EGFP-tagged condition/mKATE of EGFP untagged condition) to quantify change in degradation capacity upon adding the degron ± the ratio of the standard deviation to the mKATE mean multiplied by change in capacity. Data derive from three independent experiments each comprised of two biological repeats. A two-tailed Student T-Test was used, where P values are denoted as follows: *****<0.00005, ****<0.0005, ***<0.0005, **<0.005, *<0.05. The number of biological repeats for each sample and exact P values are reported in the Source Data File.

As previously observed for bacteria^31,33^, our expectation was that competition for degradation resources between the MODCPEST-tagged test plasmid with the PEST-tagged capacity monitor would result in coupling of the intracellular levels of the two fluorescently-tagged proteins indicating a possible bottleneck or queueing effect in the degradation pathway. This observation would be in line with the notion that the MODCPEST-tagged EGFP necessitates more protein degradation resources than its untagged counterpart, consequently reducing the availability of resources for efficient degradation of mKATE-PEST at its usual rate and thus leading to increased mKATE expression.

When MODCPEST was added to the EGFP-expressing construct, a reduction of the intracellular EGFP levels to 37% in HEK293T and 31% in CHO-K1 was observed compared to the untagged construct. This was mirrored by a 2.7- and 3.2-fold increase in mKATE from the co-transfected capacity monitor (Figure 1B).

We speculate that this restoration is not attributed to increased expression levels of the mKATE gene but rather results from reduced degradation of mKATE, a phenomenon akin to previously documented evidence in bacterial systems^31,33^. Our experiment indicates that the change in the capacity monitor’s degradation is caused by the addition of a MODCPEST signal to the competing test construct, signifying a competition for resources. The MODCPEST-tagged EGFP variant requires more protein degradation resources compared to its untagged counterpart, thereby diminishing the resources available for efficient degradation of the PEST-tagged capacity monitor at its typical rates and leading to intracellular accumulation of the mKATE protein.

### A library of degradation domains identifies specific degradation resource bottleneck

Once confirmed that we could adopt transient monitors to track competition for degradation resources in mammalian cells, we designed a suite of transient capacity monitors and test constructs that would enable us to assess and characterise the resource footprint of different degradation domains (Figure 2A). We sought to consider both N-terminal and C-terminal degrons, both stronger and weaker than MODCPEST, as per their prior characterization^17^. We thus designed a library of seven EGFP expressing constructs bearing different degrons (UbVR, 2xUbAV, UbR, UbM, MODCPEST, PEST and a composite degron with an N-terminal UbVR and a C-terminal PEST).

**Figure 2.**
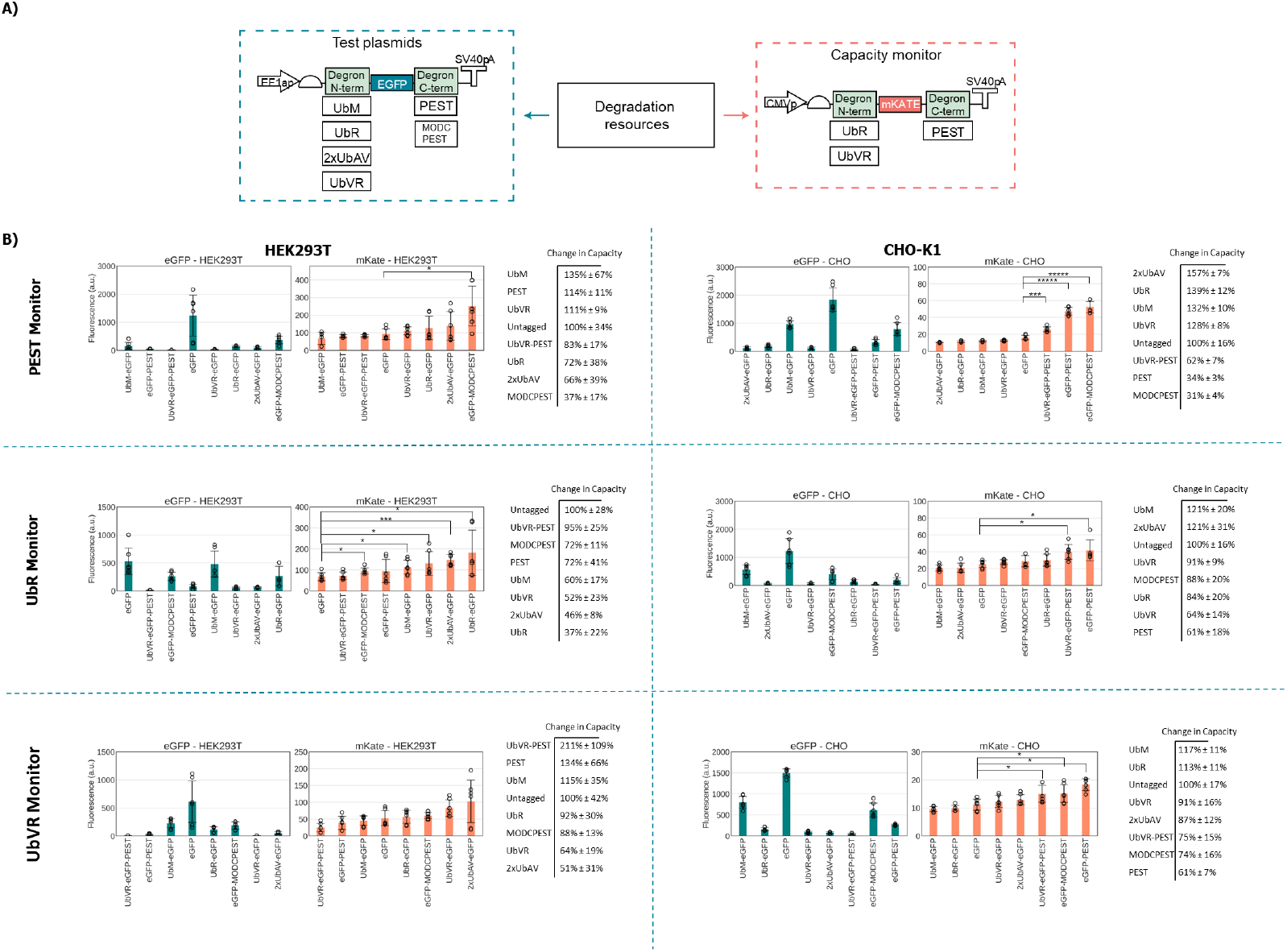
Characterizing a library of degrons with three co-transfected capacity monitors. **A)** Schematic of the genetic construct library considered. Three degradation monitors tagged with UbR, UbVR and PEST respectively were co-transfected with seven test constructs tagged with UbM, UbR, 2xUbAV, UbVR, PEST or MODCPEST. **B)** Flow cytometry data for monitor with PEST (top), UbR (middle) and UbVR (bottom), in HEK293T (left) and CHO-K1 cells (right). Test plasmid and capacity monitor reporter levels are reported as mean fluorescence (arbitrary units) ± std. Data derived from three independent experiments each with two biological repeats. A two-tailed Student T-Test was used, where P values are denoted as follows: *****<0.00005, ****<0.0005, ***<0.0005, **<0.005, *<0.05, and it was annotated only on the mKATE charts with reference to the control, EGFP versus capacity monitor, for clarity purposes. Tables report 1/(mKATE of EGFP-tagged condition/mKATE of EGFP untagged condition) to quantify change in degradation capacity upon adding the degron ± the ratio of the standard deviation to the mKATE mean multiplied by change in capacity. The number of biological repeats for each sample and exact P values are reported in the Source Data File. Tables reporting the percentage change incapacity monitor expression when constructs bear EGFP tagged for degradation compared to untagged EGFP are reported.

In order to run competition assays, we designed two additional transient capacity monitors utilising UbR and UbVR.

By co-transfecting different combinations of test constructs and capacity monitors and assessing EGFP and mKATE fluorescence by flow cytometry at 48 hours post transfection, we expected to map competition patterns for different degrons and identify degron-specific competition. To capture the impact of the cellular background on competition, we performed the analysis in both CHO-K1 and HEK293T cells.

In CHO-K1 cells, C-terminal degrons, PEST, MODCPEST, and UbVR-PEST showed a higher propensity for coupling by causing an increase in the levels of the capacity monitor, irrespective of the capacity monitors tested, up to ~3.3-fold difference (Figure 2B, C and D). In HEK293T cells, only MODCPEST induced significant coupling when co-transfected with the PEST monitor, leading to a ~2.7-fold increase in mKATE (Figure 2B). The PEST and MODCPEST degrons resulted in mild coupling with the UbR monitor both with a ~1.4-fold increase in mKATE (Figure 2C). With the N-terminal degron UbR as capacity monitor instead, it is the N-terminal capacity monitor that has shown the highest degree of coupling: UbVR with a ~1.9-fold increase, 2xUbAV with ~2.2-fold and UbR with ~2.7-fold, while with UbVR the same were ranked highest but with a much milder effect: ~1.6-fold, ~2-fold and ~1.1-fold increase in mKATE, respectively (Figure 2C and D).

Interestingly, competition for degradation resources in our experiments does not appear to be linearly correlated with the strength of the degrons employed. Therefore, context, such as position of the degron on the protein sequence and cell line, and degradation pathway differences may play a major role in the establishment of protein degradation bottlenecks.

### Inducible degradation domains enable resource-orthogonal degradation

Inducible degrons can be used to trigger protein degradation upon addition of a chemical. Their fast dynamics and ease of control through cell-permeable small molecules have led to inducible degrons being adopted as tuning dials for protein regulation both in synthetic biology and cell therapy^21–23,27^. However, given that most known degrons share part or the whole degradation pathway across organisms, the development or identification of orthogonal degrons, enabling orthogonal independent protein degradation control has become of paramount importance for their wider applicability.

We sought to test the effect of two commonly used inducible degradation domains, mAID2, an auxin-inducible degron^23^ and ecDHFR-DD, a trimethoprim (TMP) inducible degron^22^, on a co-transfected UbVR-tagged capacity monitor. Both domains were fused to the N-terminus of EGFP. to resemble UbVR, also located at the N-terminal. We reasoned that degrons derived from different organisms would be more likely to use different enzymes of the degradation pathway. For example, mAID2 requires its independent ligase to be heterologously expressed.

While both systems allow for inducible control over the stability of a protein of interest, they do have significant differences. The *Escherichia coli* dihydrofolate reductase protein (ecDHFR) is a 45-Kda degradation-prone domain stabilized through the introduction of the small molecule antibiotic TMP^22^ (Figure 3A). We cloned a plasmid containing a DHFR-EGFP fusion protein and co-transfected it alongside our UbVR-mKATE capacity monitor design.

**Figure 3.**
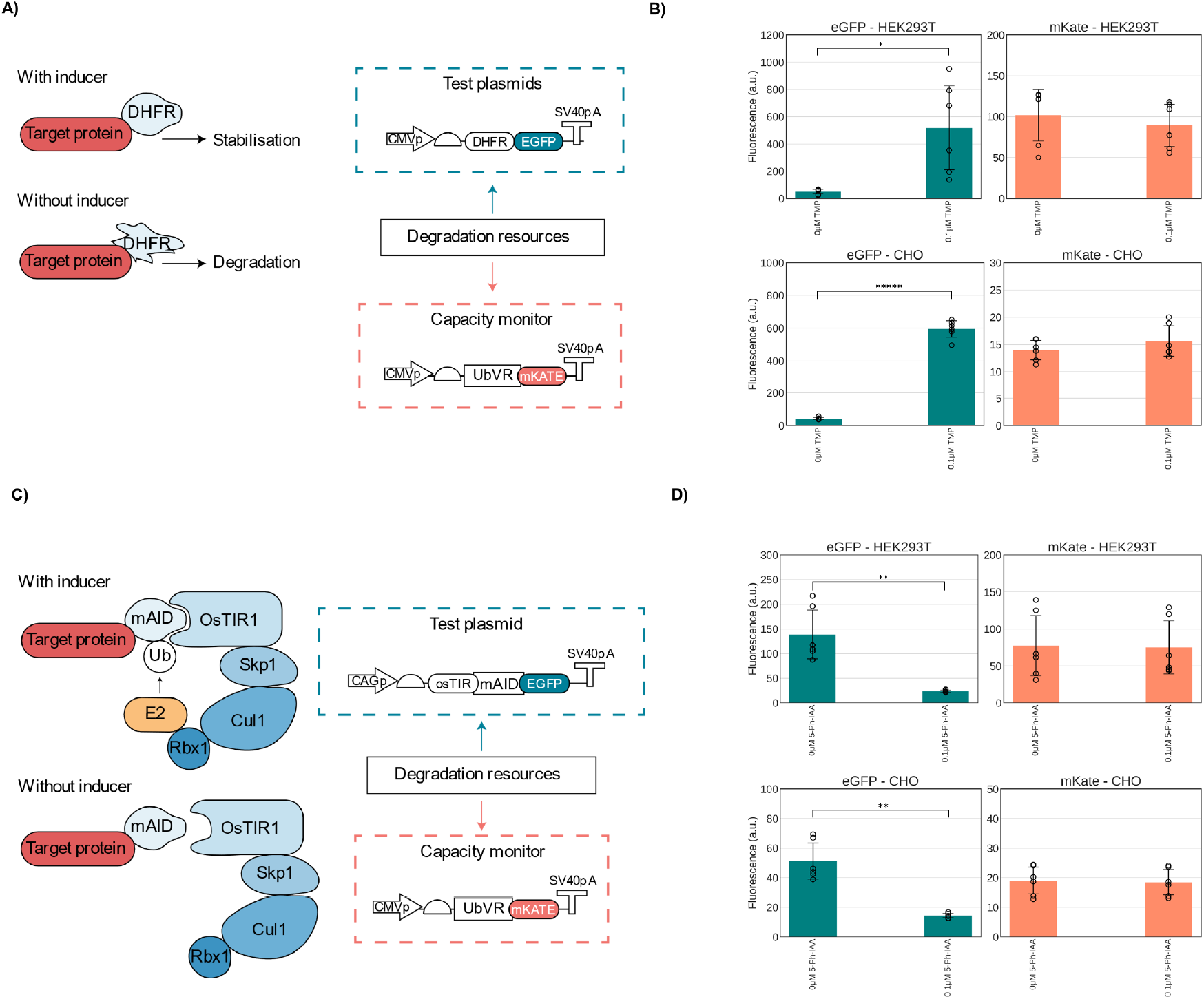
Testing bacterial and plant degrons for coupling in mammalian cells. **(A)** Schematic diagram of the TMP-inducible DHFR degron and the experimental setup. The degron is inherently destabilizing, TMP binds it and stabilizes it. These were tested against the UbVR degron and their uninduced condition was compared to the induced one. **(B)** The UbVR destabilized monitor was co-transfected in both CHO-K1 and HEK293T with an EGFP bearing the TMP degron either uninduced or induced with 0.1μM of TMP. **(C) D**iagram of the 5-Ph-IAA-inducible mAID2 degron and experimental setup. OsTIR1 forms an Skp1-Cul1-Fbox (SCF) E3 ligase complex with endogenous mammalian components. The degron is activated upon addition of 5-Ph-IAA which coordinates binding of the osTIR1 E3 ligase to the mAID2 degron. These were tested against the UbVR degron and their induced condition was compared to the uninduced one. **(D)** The UbVR destabilized monitor was co-transfected in both CHO-K1 and HEK293T with a construct encoding the OsTIR1 E3 ligase and EGFP bearing the mAID2 degron either uninduced or induced with 0.1μM of 5-Ph-IAA. Test plasmid and capacity monitor intracellular protein levels are reported as mean fluorescence (arbitrary units) ± st**d**. Data derive from three independent experiments each comprised of two biological repeats. A two-tailed Student T-Test, where P values are denoted as follows: *****<0.00005, ****<0.0005, ***<0.0005, **<0.005, *<0.05. The number of biological repeats for each sample and exact P values are reported in the Source Data File.

Addition of 0.1μM of TMP, the concentration which was previously shown to maximise stabilization in mammalian cells^22^, did not lead to a statistically significant change in intracellular mKATE levels, while intracellular EGFP levels increased up to ~14-fold (Figure 3B). The increase in EGFP suggests the correct functioning of the N-terminal DHFR domain fused to EGFP in both cell lines. Therefore, the stability of mKATE levels across conditions concomitant with a reduction in EGFP degradation in both cell lines suggest that degradation resources required for the TMP-inducible degron are at least partially orthogonal to the ones required for the N-terminal degron UbVR, similarly to previous work.

The auxin-degradation technology relies on three key components: the *Oryza Sativa* TIR1 protein F74G (OsTIR1 F74G), a 7-KDa degron known as mAID2 fused to the protein of interest, and the small-molecule auxin. When expressed in mammalian cells, OsTIR1 forms an Skp1-Cul1-Fbox (SCF) E3 ligase complex with endogenous components. In the presence of 5-Ph-IAA, this complex binds to mAID2, leading to the ubiquitination and subsequent proteasomal degradation of the protein to which it is fused (Figure 3C). This technology enables efficient protein degradation in human cells expressing OsTIR1, with a half-life of 60-65 minutes^23^.

When co-transfecting the mAID2-tagged EGFP with the UbVR-tagged capacity monitor, a decrease between 3.5 and 20-fold in EGFP levels upon the addition of 0.1μM of the inducer molecule 5-Phenyl-1H-indole-3-acetic acid (5-Ph-IAA) was observed across cell lines, while we failed to observe significant differences in mKATE (Figure 3D). Similarly to our TMP-inducible degron, these findings suggest that degradation resources required for the 5-Ph-IAA-inducible degron are at least partially orthogonal to the ones required for the N-terminal degron UbVR.

### Competition for Gene Expression and Degradation Resources Affects Genomically Integrated Genes

To investigate whether competition for degradation resources impacts the behavior of integrated synthetic cassettes, we constructed a capacity monitor cell line in HEK293T. To achieve fast, sensitive and controllable response to resource uptake from transcription to degradation, we designed the monitor to be inducible and to bear N- and C-terminal degradation tags.

We cloned a construct where, upon doxycycline induction, the TET promoter (TETp)^49^ drives the expression of three mCherry genes fused together and appended with an N-terminal and C-terminal degradation domains, UbVR and PEST, respectively. We integrated this design at the AAVS1 locus in HEK293T cells with a CRISPR-Cas9 all-in-one plasmid^50^ (Figure 4A) (see also the Methods section and Related Manuscript File 1). We selected clone 3-B7 for further experiments as it displayed correct PCR-genotyping profile and the expected induction behavior in the presence of the inducer (Figure 4B).

**Figure 4.**
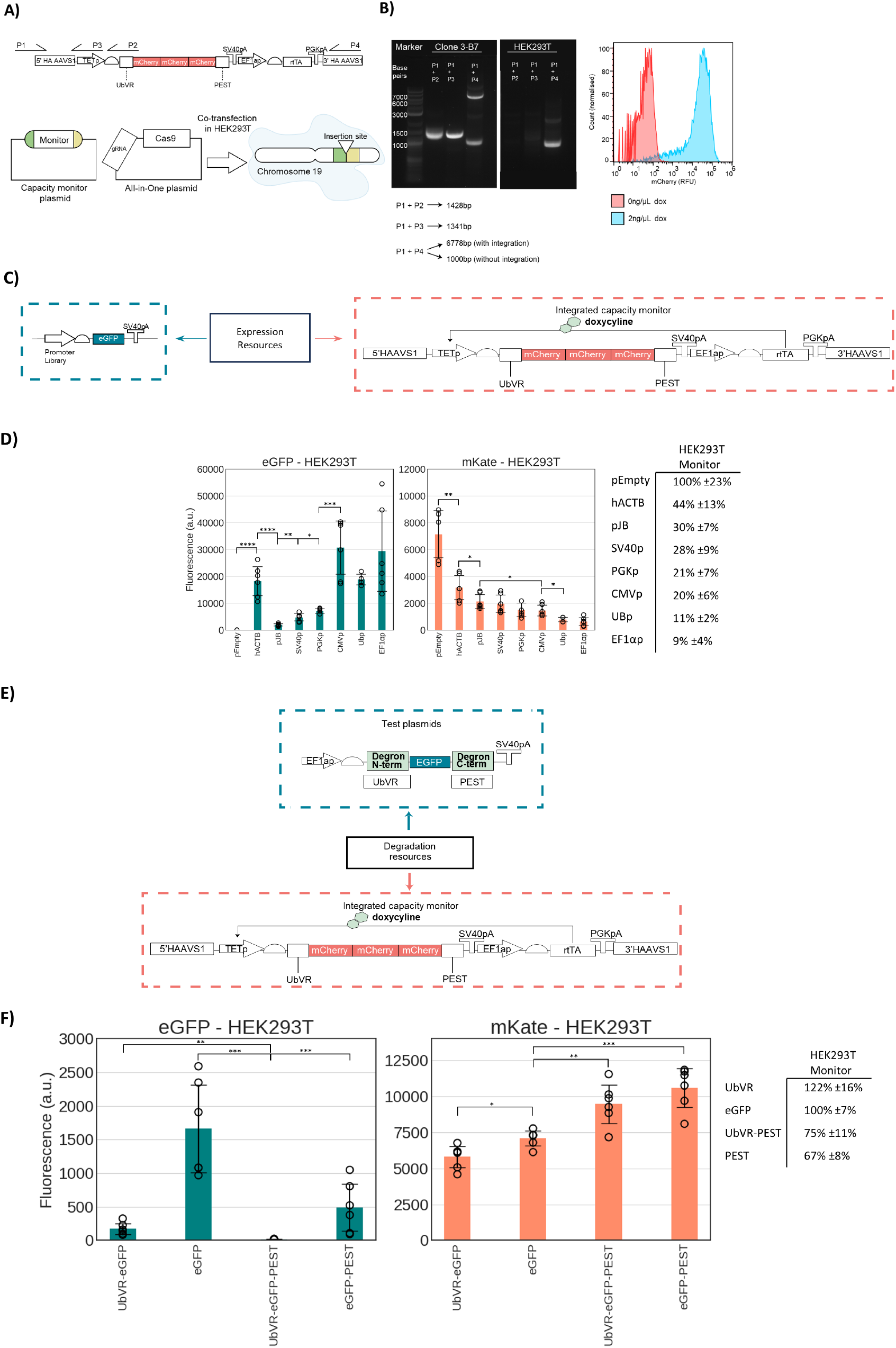
Resource competition and degradation coupling between transfected and integrated genes. **(A)** Diagram of the construction of an integrated, degraded, capacity monitor in HEK293T. The monitor is composed of three mCherry repeats flagged by both N-terminal and C-terminal degrons. It was integrated with an All-in-one CRISPR system into chromosome 19 at the AAVS1 safe harbor locus. **(B)** Integration validation via PCR and flow cytometry of the expanded PCR-verified clone. **(C)** A library of test plasmids bearing different promoters was selected to test the monitor response to competition. **(D)** EGFP construct expression and mKATE monitor expression when the promoter library is adopted. A summary table reports (mKATE of EGFP expressing condition/mKATE of pEmpty condition). **(E)** Competition for degradation resources with the integrated capacity monitor tested against EGFP bearing either UbVR, PEST or both. **(F)** Test plasmids were transfected along empty plasmids to match the total DNA transfected to the previous experiments. The table reports 1/(mKATE of EGFP-tagged condition/mKATE of EGFP untagged condition). Test plasmid and capacity monitor intracellular protein levels are reported as mean fluorescence (arbitrary units) ± std. Data derive from three independent experiments each comprised. A two-tailed Student T-Test was used, where P values are denoted as follows: *****<0.00005, ****<0.0005, ***<0.0005, **<0.005, *<0.05. The number of biological repeats for each sample and exact P values are reported in the Source Data File.

Since competition for transcriptional and translational resources by capacity monitor measures in mammalian cells has only been tested in transient systems^34–36^, we first sought to investigate if an integrated monitor can quantify competition for gene expression resources.

Previous work highlighted that transcriptional resource load plays a major role in determining the footprint of a genetic constructs on the host gene expression resources^35,36^. we started by considering monitor expression when the monitor cells are transiently transfected with a library of constructs that only differ in their EGFP promoter (Figure 4C).

The library was previously tested in Di Blasi *et al*. 2023^35^ and includes seven constitutive promoters, the two viral CMVp (Cytomegalovirus promoter) and SV40p (Simian Virus 40 promoter), and five human, pJB (a minimal promoter derived from JunB) hACTB (human β-actin), EF1αp (elongation factor 1α promoter), UBp (Ubiquitin promoter) and the PGKp (3-phosphoglycerate kinase promoter).Increasing promoter strength on the EGFP constructs yielded up to ~11-fold decrease in monitor expression, confirming that the integrated monitor is sensitive to increasing transcriptional resource uptake. A single exception was represented by hACTB, resulting in reduced monitor expression capacity by ~56% but yielding EGFP levels considerably higher than more burdensome promoters (Figure 4D). This is in line with previous work highlighting that some promoter pairs can lead to a trend different from the linear decrease in monitor expression with higher EGFP expression usually expected^35^. Growth was shown to linearly correlate with monitor expression, with decreasing cell counts observed for decreasing levels of the capacity monitor, mKATE expression (Figure S1).Once confirmed that the monitor can be adopted as proxy for competition between genomically expressed and transient genetic cassettes, we extended our analysis to degradation resources. We transfected the HEK293T capacity monitor cell line with test plasmids encoding either an untagged EGFP or an EGFP fused with UbVR, PEST or both UbVR and PEST degradation domains (Figure 4E). Remarkably, upon transfection of PEST and UbVR-PEST tagged constructs, we observed increased accumulation levels of the capacity monitor, 1.49-fold (p-value = 0.0002) and 1.33-fold (p-value = 0.005), respectively (Figure 4F). Interestingly, while the UbVR-tagged EGFP experienced more degradation than the PEST-tagged EGFP, 9.71-fold reduction (p-value = 0.0005) versus 3.38 (p-value = 0.005) compared to the untagged construct, the latter resulted in higher accumulation levels of the integrated capacity monitor, 1.49-fold (p-value = 0.0002) increase versus 1.22-fold (p-value = 0.01) decrease (Figure 4F). This observation aligns with our earlier findings regarding the competition of two C-terminal degradation tagged proteins (Figure 1B, Figure 2B). It further suggests that certain critical C-terminal or PEST-specific cellular resources may become limited when two PEST-tagged proteins are expressed simultaneously.

## Discussion

Degrons are amino acid sequences that modulate the degradation rate of the protein they are fused to, often by triggering the ubiquitin-proteasome system. They have been widely adopted in synthetic biology to tune protein degradation rate and achieve desired functionalities^8,13,17,21,27,28^. In bacteria, previous research highlighted how degradation of engineered heterologous proteins can suffer from bottlenecks derived from limitations in the cellular protein degradation machinery^30–33^. As of today, such a characterisation is still lacking for mammalian cells limiting our ability to include considerations of such limitations at the design stage and improve predictability.

Here we characterise the resource footprint of a diverse library of N-terminal and C-terminal degradation domains when degron-bearing constructs are co-transfected with a transient capacity monitor. While we observe competition for nearly all the degron-bearing constructs considered, our data highlight that in the case of C-terminal degrons such as PEST and MODCPEST, a higher degree of coupling is present, suggesting that specific bottlenecks may exist for C-terminal degradation. We speculate that this could be justified by the limited capacity of the ubiquitin-independent degradation pathway (UbInPD). The UbInPD was estimated to act on only 20% of the proteome^51^, and subsequent studies highlighted that proteins degraded in a ubiquitin-independent manner are preponderantly characterised by the presence of C-terminal degrons^52^. Furthermore, PEST-like sequences in multiple proteins have previously been shown to be degraded through the UbInPD^42–44,53,54^.

If ubiquitin was the bottleneck, we would have expected to observe competition strongly affecting all the N-terminal degrons tested which rely on ubiquitin. Further evidence in cells and mice suggests the ubiquitin pool to be highly adaptable to changes in the cell’s degradation requirements, therefore, unlikely to be a bottleneck in a healthy cell^55–57^. If the proteasome was the largest contributor to competition, we would have observed it across all the degrons tested as they require the endogenous proteasome to be degraded. However, the specific bottlenecks causing competition in the degradation pathway and the mechanistic differences between the effects of Ub-dependent and Ub-independent degrons on resource competition will need to be investigated in more detail in future studies. For instance, employing a diverse set of K48 (lysine 48) polyubiquitinated fusion proteins, which can be directly recognized and degraded by the proteasome, and observing their effect on a degradation monitor would help ascertain whether the largest bottleneck is indeed the proteasome^58^.

We then quantify competition imposed by N-terminal inducible degrons, DHFR degradation domain and mAID2, respectively and observe no significant coupling with the capacity monitor. We speculate that mAID2 may rely on partially orthogonal, plant-derived, E3 ligase complex, the Skp1-Cul1-osTIR1^23,59^. While ecDHFR is a bacterial unstable domain, its degradation pathway in mammalian cells has not been fully elucidated at the molecular level^22^. Overall, our experiments provide evidence that, when proteins subject to N-terminal degradation are present within an engineered synthetic circuit, systems like these inducible degrons could be adopted offering the advantage of reduced competition and possible related circuit mis-functioning.

To investigate the impact of transiently expressed cassettes on genomically integrated genes, we then construct a capacity monitor in HEK293T which we first test for expression competition with a promoter library previously tested with transient monitors. The promoter library confirmed that competition for transcriptional resources, can be quantified by means of an integrated monitor showing an inverse linear correlation between expression levels of the transient construct and the integrated monitor with hACTB as the only exception.

Moreover, the integrated capacity monitor, when tested with plasmids from our constitutive degrons library, confirmed coupling of intracellular resources between transient and integrated genes in the degradation pathway too. Similarly to the transient system, in the integrated monitor, C-terminal or PEST-like degrons appear to compete for a smaller pool of resources even though they exhibit degradation levels comparable to N-terminal degrons. Future fundamental research should focus on understanding the cellular repercussions of competition for degradation, as we show it can affect genomically integrated genes and targeted degradation therapeutics are entering the clinic.

The integrated capacity monitor presented here functions as a proxy measure of expression resource availability in HEK293T and provides an example of how integrated monitors could be leveraged in mammalian cell engineering. Our integrated monitor, while it is not specific like qPCR or RNA-seq in pinpointing changes in expression levels, offers a live dynamic proxy measurement of expression and degradation competition and its impact on the cell, offering a complementary approach to cell growth for burden assessment. Further research will be needed to elucidate the effect of this competition on a broader range of genomic genes and across different cell lines^58^.

Overall, in the future, bottlenecks in protein degradation could be leveraged in synthetic biology to improve context-aware design of gene circuits exploiting degrons or to induce competition for degradation as a mean of synthetic regulation. For therapeutic purposes, competition for degradation could be used to control specific protein degradation pathways and to stabilise cellular proteins whose dysregulated degradation contributes to human diseases^60–62^ surpassing the toxicity challenges of current proteasomal inhibitors^63–65^ or to optimise the multiplexing of PROTACs or molecular glues because of their reliance on the endogenous degradation pathways^66–68^.

Our findings underscore the importance of taking a more holistic approach when considering the impact of competition for gene expression resources in mammalian cells, including consideration for additional regulatory pathways such as protein degradation and secretion resources. Such a comprehensive approach is critical for unraveling the intricacies of cellular regulation when disrupted by expression of heterologous genes and could have broad-reaching implications for the fields of biotechnology and synthetic biology.

## Methods

### Plasmid Design and Preparation

The plasmids adopted in this study were built by PCR-based cloning. Starting from donor plasmids and a template plasmid, we added the degradation tags at the C and N terminus of the fluorescent protein. Fragments were isolated by gel electrophoresis (Agarose, Sigma-Aldrich, Gel Loading Dye, Purple(6x), NEB) and purified using a Qiagen extraction kit as per the manufacturer’s instructions (QIAquick Gel Extraction Kit, Qiagen). A complete list of plasmids included in this study can be found in Related Manuscript File 2. Plasmid maps annotated for specific parts and sequences used in this paper can be found in the Source Data File. Primers used in this study can be found in Related Manuscript File 3.

Templates and inserts were restricted and ligated with the EMMA protocol^73^ using Esp3I (Esp3I, Thermo Scientific) and T4 DNA ligase (T4 Ligase, NEB). Ligations were transformed in chemically competent *E. coli* DH5α by heat shock at 42°C for 30 seconds and 45 minutes outgrowth in 1mL Luria Broth (LB Broth, Sigma-Aldrich). 100μL were then plated on agar plates supplemented with 100μg/mL ampicillin (Ampicillin, Sigma-Aldrich). The antibiotic-resistant colonies were then inoculated in 5mL of Luria Broth the following day and incubated overnight at 37°C. The culture was then midi prepped (CompactPrep Plasmid Midi Kit, Qiagen) according to the manufacturer’s protocol the following day to extract the plasmid DNA, ready to be diluted for transfection.

### Mammalian Cell Culture and Transfection

HEK293T cells (HEK293T, ATCC) were cultured in T75 flasks (Nunc™ EasYFlask™ Cell Culture Flasks, Thermo Scientific) using DMEM (Dulbecco’s modified Eagle Medium) supplemented with 1mM sodium pyruvate, 1x GlutaMaxTM, 0.4mM phenol red, and 25mM glucose (DMEM, high glucose, GlutaMAX™ Supplement, pyruvate, Gibco), along with 10% FBS (Fetal Bovine Serum, Gibco). Cells were kept in an incubator at 37°C and 5% CO2 and were passaged upon reaching 70% confluency. 1mL of trypsin-EDTA (0.5%, no phenol red, Gibco) was used for cell detachment during passages, and 1mL of PBS (phosphate buffered saline, Merck) was used for cell washing.

CHO-K1s were cultured in MEMa (Minimum Essential Medium Eagle, Sigma) supplemented with 10% FBS (Fetal Bovine Serum, Gibco), 1% non-essential amino acids (MEM Non-Essential Amino Acids Solution (100X), ThermoFisher), and 1% L-glutamine (L-Glutamine (200 mM), ThermoFisher) in T75 flasks (Nunc™ EasYFlask™ Cell Culture Flasks, Thermo Scientific).

HEK293T cells were seeded 1 day before transfection at 100,000 cells/well in 24-well plates (Multiwell cell culture plates, VWR) for the transient transfections. Details on the transfection mixes can be found in Related Manuscript File 4. The complete mix was incubated for 30 minutes pre-transfection and 2 days after.

CHO-K1 cells were seeded for transfection at 8*10^4^cells/well in 24w plates one day before transfection. Details on the transfection mixes can be found in Related Manuscript File 4. The complete mix was incubated for 25 minutes at room temperature pre-transfection and 2 days after.

2ng/μL of doxycycline (doxycycline, Sigma Aldrich) was added to the wells containing cells transfected with plasmids with a dox-inducible promoter for HEK293Ts on day 1. For both HEK293Ts and CHO-K1s 0.1μM 5-Ph-IAA (5-phenylalanine-indole acetic-acid) (5-Ph-IAA, Cambridge Bioscience) to the ones with an mAID2 tag and 0.1μM TMP (Trimethoprim Ready Made Solution, Sigma Aldrich) to the ones with an ecDHFR tag right after transfection or at several time points.

### Stable cell line development

This study’s CRISPR integrations involved integrative payloads expressing a fluorescence gene. This facilitated the selection of the integrated population and allowed easy assessment of the stability of the selected clones. All the integrative payloads in this study have been integrated into AAVS1 in HEK293T (described in Related Manuscript File 1), using the following gRNA sequence, which was already experimentally validated^50^:

5’-GGGGCCACTAGGGACAGGAT-3’.

For CRISPR-based integrations, 2*10^5^ HEK293T cells were seeded in 24-well plates one day before transfection. The day after, 0.06pmol of donor backbone and 0.04pmol of All-in-One backbone were co-transfected in HEK293T using FuGENEHD (FuGENE® HD Transfection Reagent, Promega) as transfection reagent at a ratio of 3:1 FuGENEHD (μL): transfected DNA (μg). 24 hours after transfection, the spent media was changed with fresh culture media and cells were cultured for 10 days at least. Cells were detached, and the positive fluorescent population for our marker was single-cell sorted into individual wells of a 96-well plate.

The resulting single-cell clones were cultured for three-to-four weeks in conditioned medium (i.e., filtered spent media collected from a confluent flask) at the ratio of 1:3 to fresh medium with 25% FBS and 1% penicillin-streptomycin (Penicillin-Streptomycin, Sigma Aldrich). Confluent wells were then assessed with fluorescence microscopes, and the ones displaying post-integration homogeneous fluorescence were sorted and expanded.

For cell sorting experiments during cell line development, cells were detached, centrifuged for 5 minutes at 3000rpm, washed in DPBS, centrifuged again for 5 minutes, and resuspended in DPBS at a final concentration of 10^7^ cells/mL. Cells were kept on ice and filtered in test tubes with cell strainer caps to disrupt cell clumps. Sorting was carried out at 4°C using a BD FACSAria Fusion (BD FACSAria Fusion, BD Biosciences) by specialized facility staff. Purity checks on the sorted population were carried out whenever possible.

As soon as they reached confluency in a 24-well plate format, the genomic DNA of the expanded clones was extracted with a kit according to the manufacturer’s protocol (Nucleospin Tissue, Macherey-Nagel) to perform PCR genotyping. Correct clones were expanded further, aliquoted, and frozen in culture media with 10% DMSO (Dimethyl Sulfoxide, Merck).

### Detection of Intracellular Fluorescence via Flow Cytometry

At 48 hours post-transfection, cells were detached by washing with 400μL/well of PBS. The cell suspension was transferred to microcentrifuge tubes and centrifuged at 1500 rpm for 5 minutes. The resulting cell pellet was resuspended in 300μL of PBS and transferred into test tubes with cell strainer caps using a filter (Corning™ Falcon™ Round-Bottom Polystyrene Test Tubes with Cell Strainer Snap Cap, 5mL, FisherScientific). For analysis, 50μL of each sample was loaded onto an Attune NxT (Attune NxT Flow Cytometer, Thermofisher) to collect approximately 10,000 singlet cell events. The EGFP fluorescence was excited with a 488nm laser and detected using a 530/30nm bandpass filter, the mKATE fluorescence was excited with a 561nm laser and detected using a 620/15nm bandpass filter.

Flow cytometry data analysis was performed using the FlowJo software (FlowJo™, BD Life Sciences). Live cells were gated based on the FSC/SSC dot plots to exclude debris with low side and forward scatter. Single cells were gated based on the FSC-H/FSC-A plot. Then, the levels of EGFP and mKATE were determined by calculating the geometric mean of the entire population. Compensation of the red and green signals was automatically performed in FlowJo using a green control expressing only EGFP and a red control expressing mKATE. Further details on the gating strategy can be found in Supplementary Note 1 and Figure S2. Further detail on the processing of flow cytometry data can be found in the Source Data File.

### Live Cell Counting Assay

For live cell counting experiments, transfected cells in 24 well plates were washed in DPBS detached in 100μL of 1% trypsin-EDTA, and 200μL of DMEM + 10% FBS was added to a total volume of 300μL. Lastly, 2.5μL of solution 18 (Solution 18, Chemometec) was added to 50μL of cell suspension and live cells were counted using the Nucleocounter (Nucleocounter NC250, Chemometec). Cell counts can be found in the Source Data File.

## Acknowledgements

The authors would like to thank Tom Ellis, Mustafa Khammash and Velia Siciliano for early discussion on the project; Martin Fussenegger for sharing plasmid pCHX116 (2xUbAV), pCHX204 (UbM), pCHX247 (UbR) and pCHX155 (MODCPEST) with the degrons adopted in this paper; Ervin Welker for sharing plasmid pSC1-DD (EGFP-ecDFHR); Karen Polizzi for constructive feedback on the manuscript. The authors acknowledge the support of the EPSRC Centre for Doctoral Training in BioDesign Engineering (EP/S022856/1) (to JG, CK and FC). R.D.B. was supported by the Imperial College Chemical Engineering PhD scholarship.

## Author contributions

All authors conceived the research and designed the experiments. JG and RDB performed the experiments. All authors wrote and edited the manuscript.

