## Supplementary Information for "Degradation Bottlenecks and Resource Competition in Transiently and Stably Engineered Mammalian Cells"

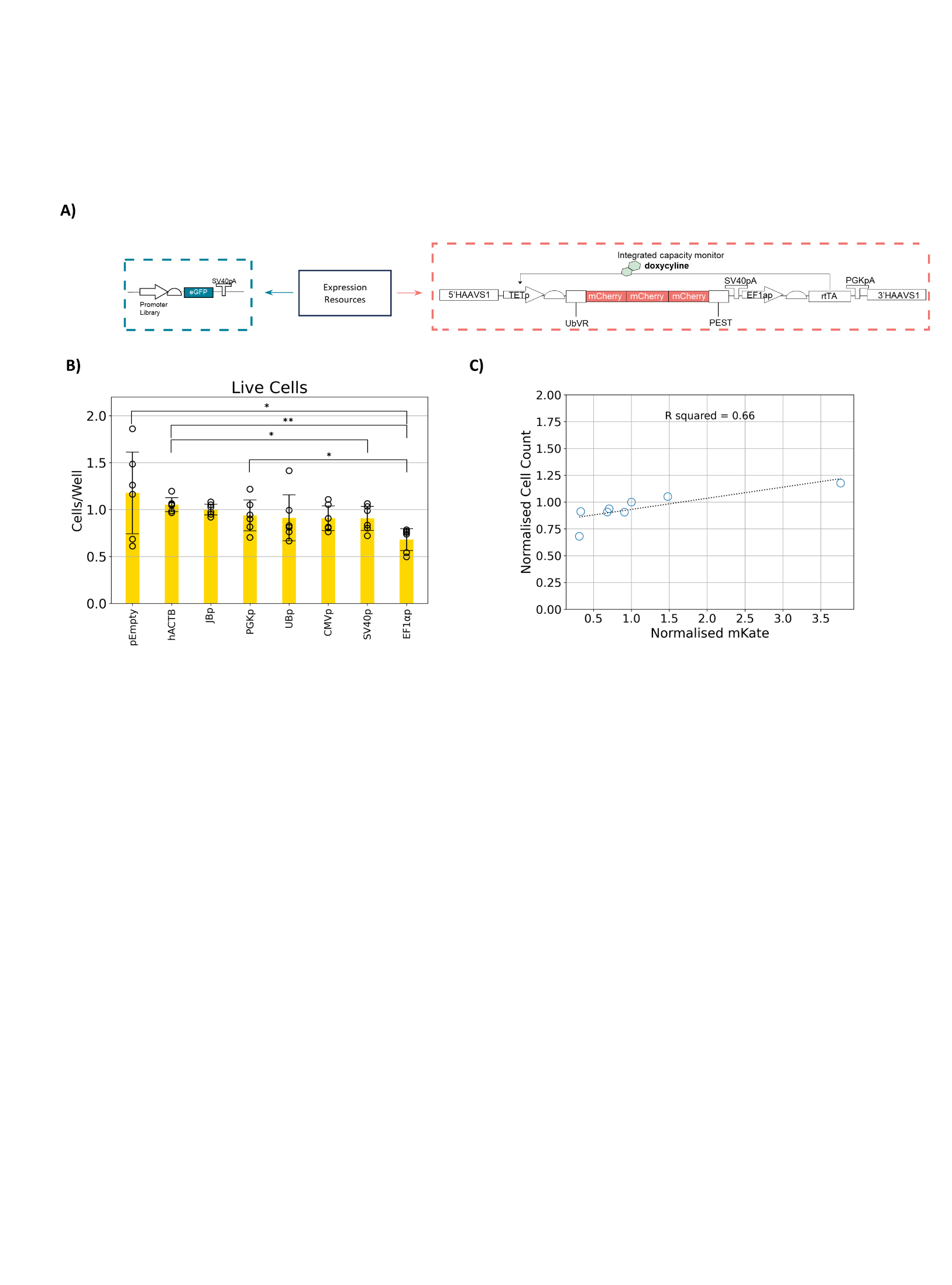

**Figure S1**. **A)** A library of test plasmids bearing different promoters was selected to test the monitor response to competition. **B)** Cell count data after two days of expression of the test plasmid versus the induced integrated capacity monitor. In these analyses, cell counts are reported as normalized mean cell count (fold-change) ±standard deviation. **C)** Scatter plot of normalised mKATE and normalised cell count with linear regression represented as a dotted line and annotated R^2^. The data presented are derived from three independent experiments each comprised of two biological repeats. Statistical significance was determined using a two-tailed Student T-Test, where P values are denoted as follows: *****<0.00005, ****<0.0005, ***<0.0005, **<0.005, *<0.05. The number of biological repeats for each sample, the normalization, and exact P values are reported in the Source Data File.

**Supplementary Note 1**

**Transfection Flow Cytometry Data Gating Strategies in the Context Resource Competition**

Resource competition studies are performed via transfection/transformation of one plasmid into a cell carrying a “capacity monitor”, a fluorescent reporter resource-coupled to the plasmid, or multiple plasmids when the “capacity monitor” is not genomically integrated. The analysis of transfection data is usually either carried in bulk via fluorescence or luminescence in a plate reader, via mRNA levels with qPCR and RNAseq or at the single cell level with flow cytometry or imaging techniques. In our article, we adopted flow cytometry given its sensibility and the granularity of the output. Flow cytometry allows to gate populations of cells, excluding ones that might not be of interest for the analysis, or might confound it, such as debris, dead cells, doublets, untransfected. Outlined here is the choice of our gating strategy:

1. Gating out debris and dead cells via an FSC/SSC gate.

2. Gating out of doublets via an FSC-A/FSC-H gate.

3. Statistical computation of geometric mean fluorescence of gated cells.

This gating strategy tends to underestimate competition effects and introduces variability due to the unpredictable variance in transfection efficiency. We consider the strategy to be high accuracy but low precision. However, alternative strategies based on gating fluorescent markers are high precision but risk low accuracy in a resource competition setting.

Gating strategies based on fluorescent markers employ either a gate on a transfection marker, a third transfected fluorescent protein, or on a transfected fluorescent protein which is part of the experiment. Both strategies are subject to competition for expression resources by the other expression cassettes, thereby inducing a condition-dependent bias in the data:

- If the transfection marker is part of an experiment with high competition for expression resources, it might be suppressed to a level where the cell population can be mistaken for untransfected cells.
- In another possible instance, fluorescent proteins being measured as part of the experiment are being used to gate out untransfected cells, however, in different conditions, different competing constructs, their levels might drastically increase or decrease, over and underestimating the results on a condition-by-condition basis.

In competition for expression resources, we might see underestimation of the transfected population in conditions of high competition as these will lower expression levels of the monitor, while in competition for degradation experiments the opposite will happen as higher degradation competition leads to an increase in undegraded, therefore fluorescent, monitor levels.

Methods derived from gating on a transfection marker or on a fluorescent protein part of the experiment, such as gating on a fixed percentile of the markers are still affected by resource competition effects and we deem them high precision, low accuracy.

We show how gating for fluorescent transfected proteins to remove untransfected cells can mislead towards the false identification of a significant difference between the capacity monitor competing with an induced degron (Figure S2A). The histograms overlaid to the flow cytometry dot plot highlight the difference between uninduced and induced ecDHFR degron only in the eGFP channel, ~10-fold decrease (Figure S2B, C). However, in the bar plots representing the geometric mean of the gated version of the data a ~2-fold difference appears in the mKATE between the uninduced and induced condition which is not reflected in the raw data (Figure S2C).

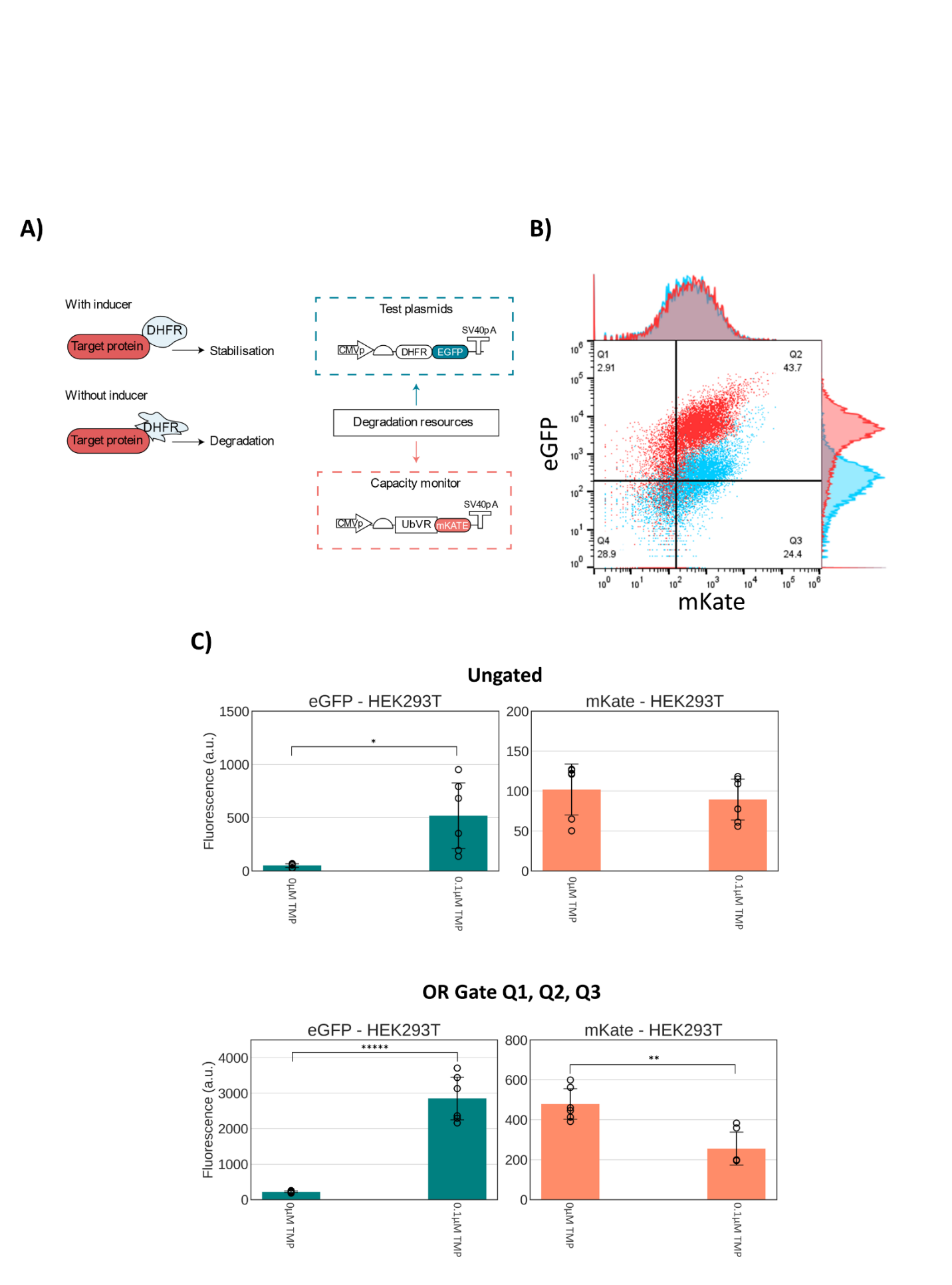

**Figure S2. A)** A diagram of the representative experiment on which we tested the two different gating strategies. **B)** A dot plot with overlaid histograms displaying in blue an uninduced ecDHFR-EGFP competing with a capacity monitor UbVR-mKATE, and in red the induced version with 0.1µM of TMP. **C)** Bar plots representing the geometric mean of flow cytometry data for EGFP and mKATE. In these analyses, test plasmid and capacity monitor intracellular protein levels are reported as mean fluorescence (arbitrary units) ± standard deviation. The data presented are derived from three independent experiments each comprised of two biological repeats. Statistical significance was determined using a two-tailed Student T-Test, where P values are denoted as follows: *****<0.00005, ****<0.0005, ***<0.0005, **<0.005, *<0.05. The number of biological repeats for each sample and exact P values are reported in the Source Data File.
